## Supplementary Information for "Computational design of dynamic receptor—peptide signaling complexes applied to chemotaxis"


**Authors**

Jefferson RE[1], Oggier A[1], Füglistaler A[1], Camviel N[2], Hijazi M[1], Rico Villarreal A[1], Arber C[2], Barth P[1, §]

**Affiliation**

1: Interfaculty Institute of Bioengineering, École Polytechnique Fédérale de Lausanne, Lausanne, CH-1015, Switzerland and Ludwig Institute for Cancer Research Lausanne, Switzerland

2: Department of oncology UNIL-CHUV, University Hospital Lausanne (CHUV), University of Lausanne (UNIL) and Ludwig Institute for Cancer Research Lausanne, Switzerland

§ Correspondences should be addressed to:

Patrick Barth, Interfaculty Institute of Bioengineering, École Polytechnique Fédérale de Lausanne, Lausanne, CH-1015, Switzerland,

**Supplementary Figure 1. Computationally-guided point mutant library activity screen.** (**A**) Receptor-mediated Gi activation normalized to cell surface expression for point mutations (x-axis) designed at different receptor positions in the binding pocket (y-axis). Untested substitutions are marked with an X. Inactive substitutions are gray. (**B**) Calcium mobilization activity for point mutants that displayed higher than WT normalized activity in the initial screen (s.d., n=3 technical replicates).

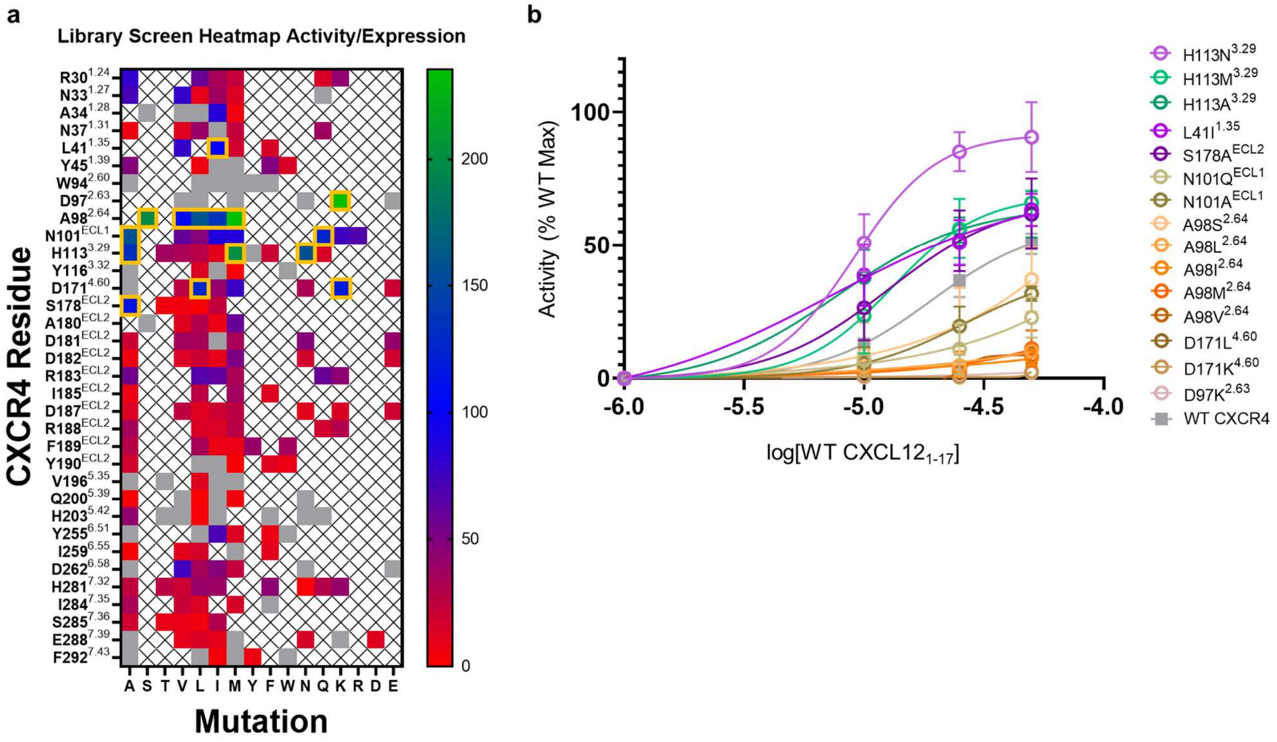

**Supplementary Figure 2. Surface expression of receptor variants.** (**A**) Surface expression of receptors in HEK 293T cells measured by ELISA (s.d., Mock: n=32 independent experiments, WT CXCR4: n=37 independent experiments, Csel1: n=5 independent experiments, Csel2: n=11 independent experiments, Cdyn: n=6 independent experiments, Csedy: n=5 independent experiments). (**B**) Surface expression of receptors in transduced T cells measured by flow cytometry (s.d., Non-transduced: n=6 biologically independent samples, WT CXCR4: n=5 biologically independent samples, Csel2: n=6 biologically independent samples, Cdyn: n=5 biologically independent samples, Csedy: n=5 biologically independent samples). Individual measurements for each T cell donor plotted.

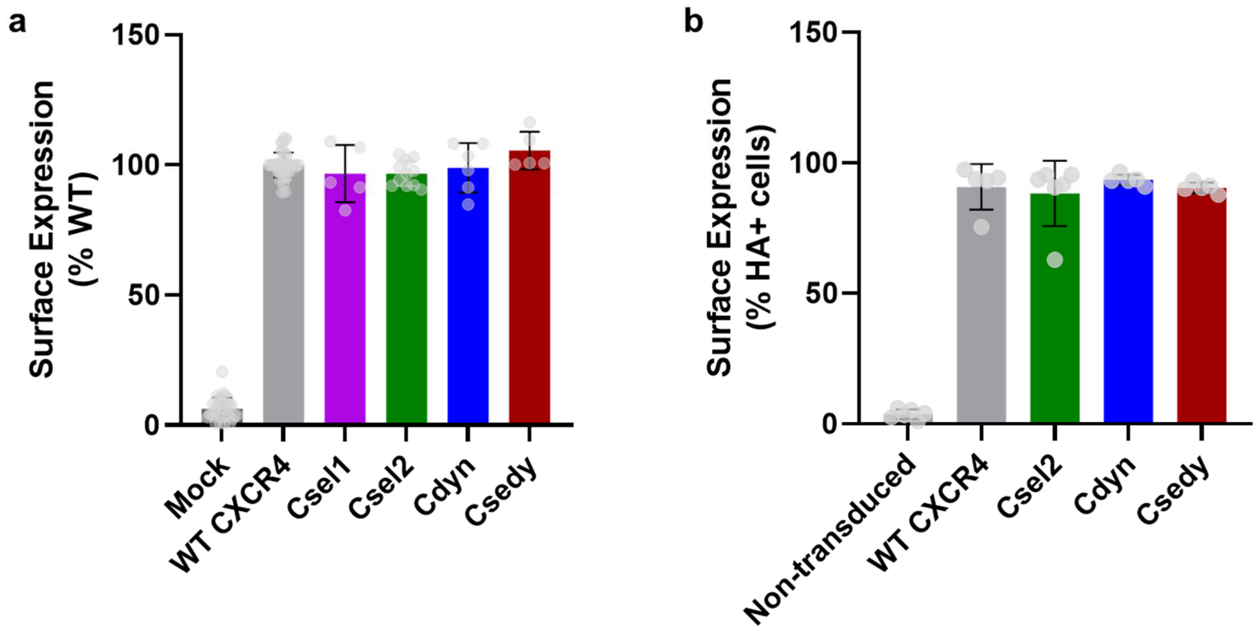

**Supplementary Figure 3. Activation landscapes of molecular dynamics simulations.** Landscapes representing the density of frames in the activation conformational space described by interhelical distances of TM3-6 and either TM3-7 on the intracellular side of the receptor (left) or the RMSD of the NPxxY motif from the inactive reference structure (right) for (**A,B**) CXCR4 WT:WT, (**C,D**) Csel2:Y7L, (**E,F**) Cdyn:V3Y, and (**G,H**) Csedy:V3Y-Y7L. Distances were calculated between alpha carbons of residues R3.50, K6.30, and Y7.53 for TM3, TM6, and TM7 respectively. Landscapes were calculated in a fashion similar to 2D potential of mean force, with densities defined by kernel density estimates with gaussian functions. The color bars are in arbitrary units, where blue (lowest quantity) represents the highest density of frames, and yellow (highest quantity) represents the edge of the populated space. Experimental structures of inactive CXCR4 (PDB ID: 4RWS) and active US28, CXCR2, CCR5 (PDB IDs: 4XT1, 6LFO, and 7F1R, respectively) chemokine receptor structures are shown for reference.

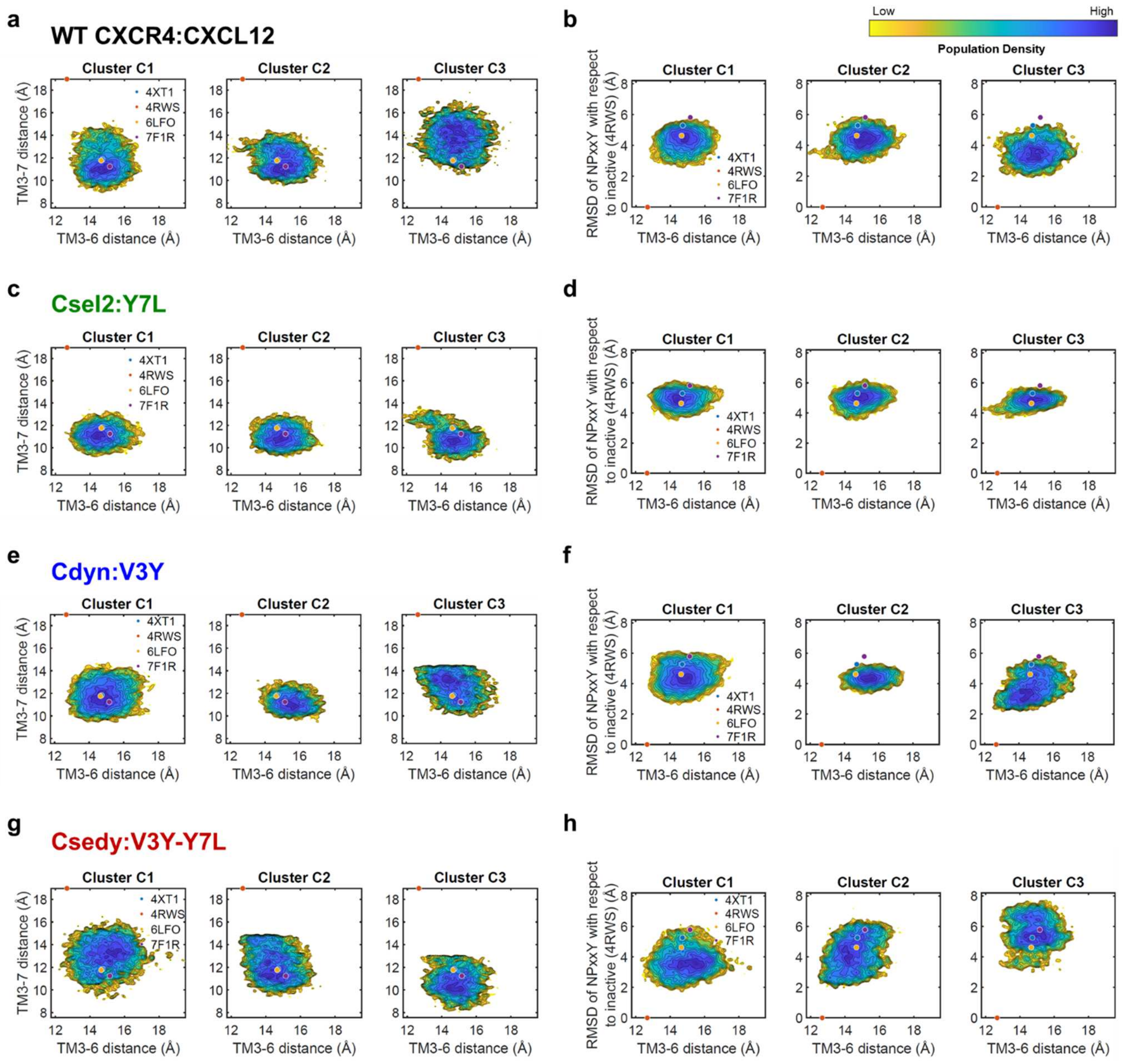
a
WT CXCR4:CXCL12
b
Low
High
Population Density
c
Csel2:Y7L
d
e
Cdyn:V3Y
f
g
Csedy:V3Y-Y7L
h
Cluster C1
Cluster C2
Cluster C3
4XT1
4RWS
6LFO
7F1R
TM3-7 distance (Å)
TM3-6 distance (Å)
RMSD of NPxxY with respect to inactive (4RWS) (Å)

**Supplementary Figure 4**: **Convergence of 1st and 2nd order entropies for WT and CAPSen receptor—peptide complexes.** Convergence plots of 1st order (blue, left Y-axis) and 2nd order (red-brown, right Y-axis) entropies as a function of number of frames in every cluster for (**A**) CXCR4 WT:WT, (**B**) Csel2:Y7L, (**C**) Cdyn:V3Y, and (**D**) Csedy:V3Y-Y7L. 1st order entropy is defined as the sum of marginal entropies of all individual dihedrals (*φ, ψ*, and *X*'s) and 2nd order entropy is the sum of joint entropy of dihedral pairs formed by the top 300 dihedrals with the highest summed MI. 1st order entropies converge much faster than 2nd order ones, so we consider 2nd order entropies for the convergence criterion. Entropies are considered converged when there is less than 10% variability from the final entropy value over the last 500 frames (50 ns) considered. The 10% limit is represented by the dashed horizontal red line. The final entropy value is marked by the solid horizontal red line. The last 500 frames are marked with the vertical dotted magenta line.

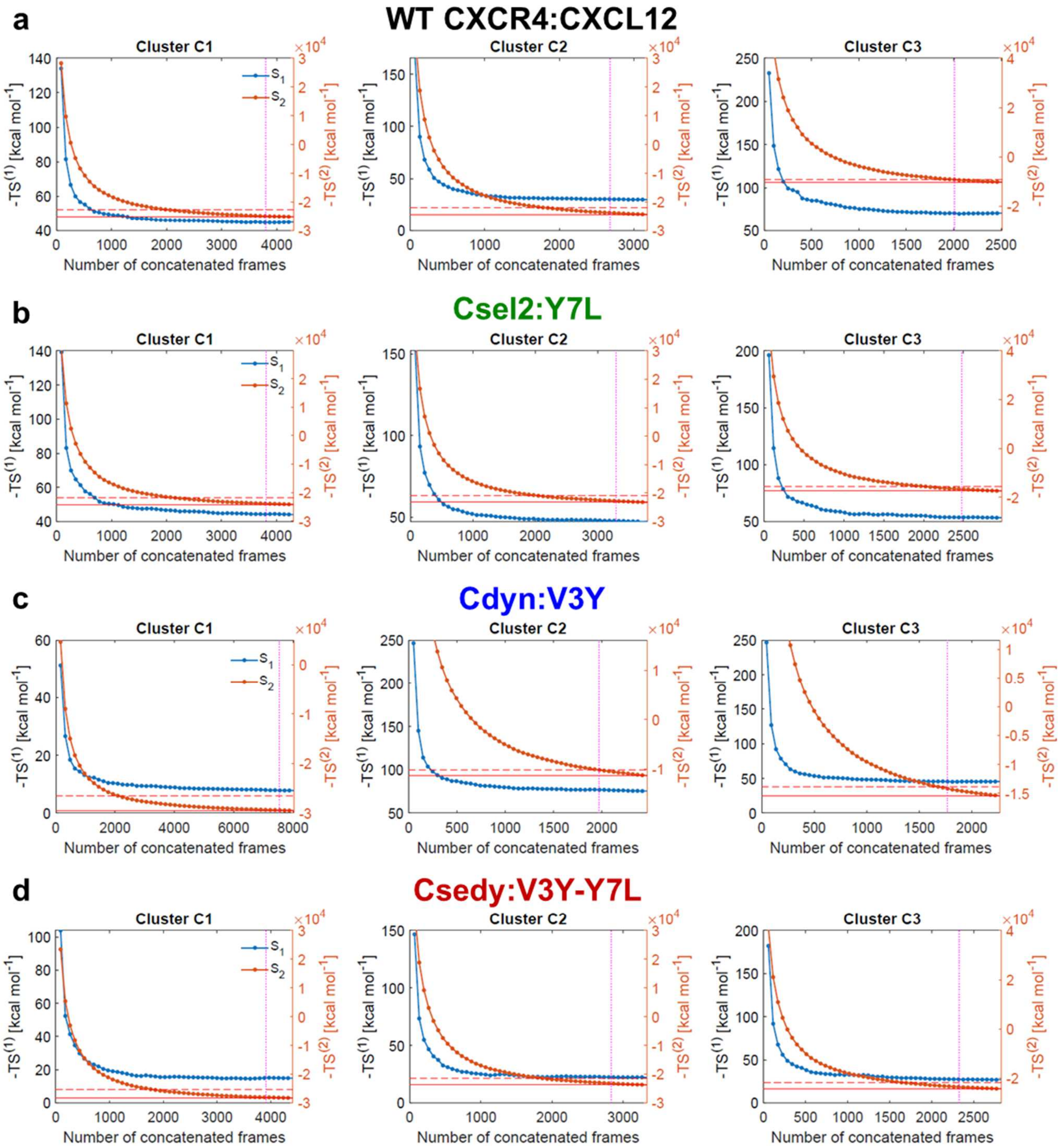
a
WT CXCR4:CXCL12
Cluster C1
Cluster C2
Cluster C3
S1
S2
-TS(1) [kcal mol-1]
-TS(2) [kcal mol-1]
×10^4
Number of concatenated frames
b
Csel2:Y7L
Cluster C1
Cluster C2
Cluster C3
S1
S2
-TS(1) [kcal mol-1]
-TS(2) [kcal mol-1]
×10^4
Number of concatenated frames
c
Cdyn:V3Y
Cluster C1
Cluster C2
Cluster C3
S1
S2
-TS(1) [kcal mol-1]
-TS(2) [kcal mol-1]
×10^4
Number of concatenated frames
d
Csedy:V3Y-Y7L
Cluster C1
Cluster C2
Cluster C3
S1
S2
-TS(1) [kcal mol-1]
-TS(2) [kcal mol-1]
×10^4
Number of concatenated frames

**Supplementary Table 1. Conformational dynamics and structural characterization of clustered substates of WT and CAPSen designs from molecular dynamics simulations.** Pathway density distribution across substates selected as input for AlloDy. Occupancy calculated from total simulated frames. Intra-cluster RMSD calculated from receptor and peptide (Complex) and contacting residues across all variants including peptide (Pocket). Contact frequency thresholds used to define binding and allosteric contacts normalized to intra-cluster complex RMSD of each substate.

| Variant | Path Density (% total) | Occupancy (% total) | Intra-cluster RMSD (Å) | |
|---|---|---|---|---|
| Substate | | | Complex | Pocket |
| **WT** | | | | |
| C1 | 51.3 | 29.6 | 2.43 | 2.54 |
| C2 | 31.9 | 21.9 | 2.55 | 2.63 |
| C3 | 16.8 | 17.3 | 2.32 | 2.30 |
| **Csel2** | | | | |
| C1 | 30.6 | 34.5 | 2.14 | 1.99 |
| C2 | 31.7 | 30.4 | 2.07 | 2.09 |
| C3 | 37.7 | 23.8 | 2.18 | 2.08 |
| **Cdyn** | | | | |
| C1 | 51.3 | 51.9 | 3.01 | 3.61 |
| C2 | 13.4 | 15.9 | 3.01 | 3.47 |
| C3 | 35.2 | 14.6 | 2.69 | 3.31 |
| **Csedy** | | | | |
| C1 | 42.8 | 30.4 | 2.51 | 2.76 |
| C2 | 18.0 | 23.0 | 2.47 | 3.04 |
| C3 | 39.1 | 19.5 | 2.48 | 2.68 |

**Supplementary Figure 5. Conformational flexibility of the ligand in the CCR5-RANTES complex.** Population density and cluster centers in PC space. Inter-cluster RMSD of the most populated ligand conformations shown. The highest density areas are colored yellow, while the lowest density areas are blue. The first 2 PCs explain 53.66% and 19.09% of the variability in the data respectively. Total simulation time is 1500 ns.

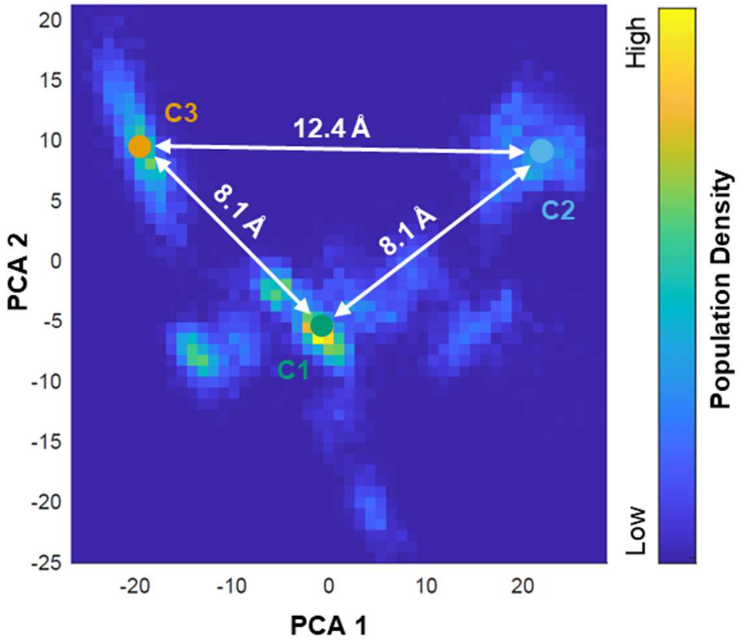

**Supplementary Figure 6. Schematic of the steps performed by AlloDy to extract correlated motions from simulations and cluster them into allosteric pipelines.** Ligand binding poses are clustered from molecular dynamics simulations using principal component analysis. A particular ligand pose is highlighted in the blue box with contact mapping to the right. Then dihedrals are extracted for every cluster separately. For every cluster, mutual information (MI) is calculated with finite size corrections and statistically filtered to remove uncoupled pairs (black) and include highly coupled pairs (red) before being summed over residue pairs. Pathways are then constructed using a shortest distance algorithm between residue pairs (red box) that have significant MI and are more than 10 Å apart stemming from ligand contacts (blue box). Constructed pathways are clustered into allosteric pipelines.

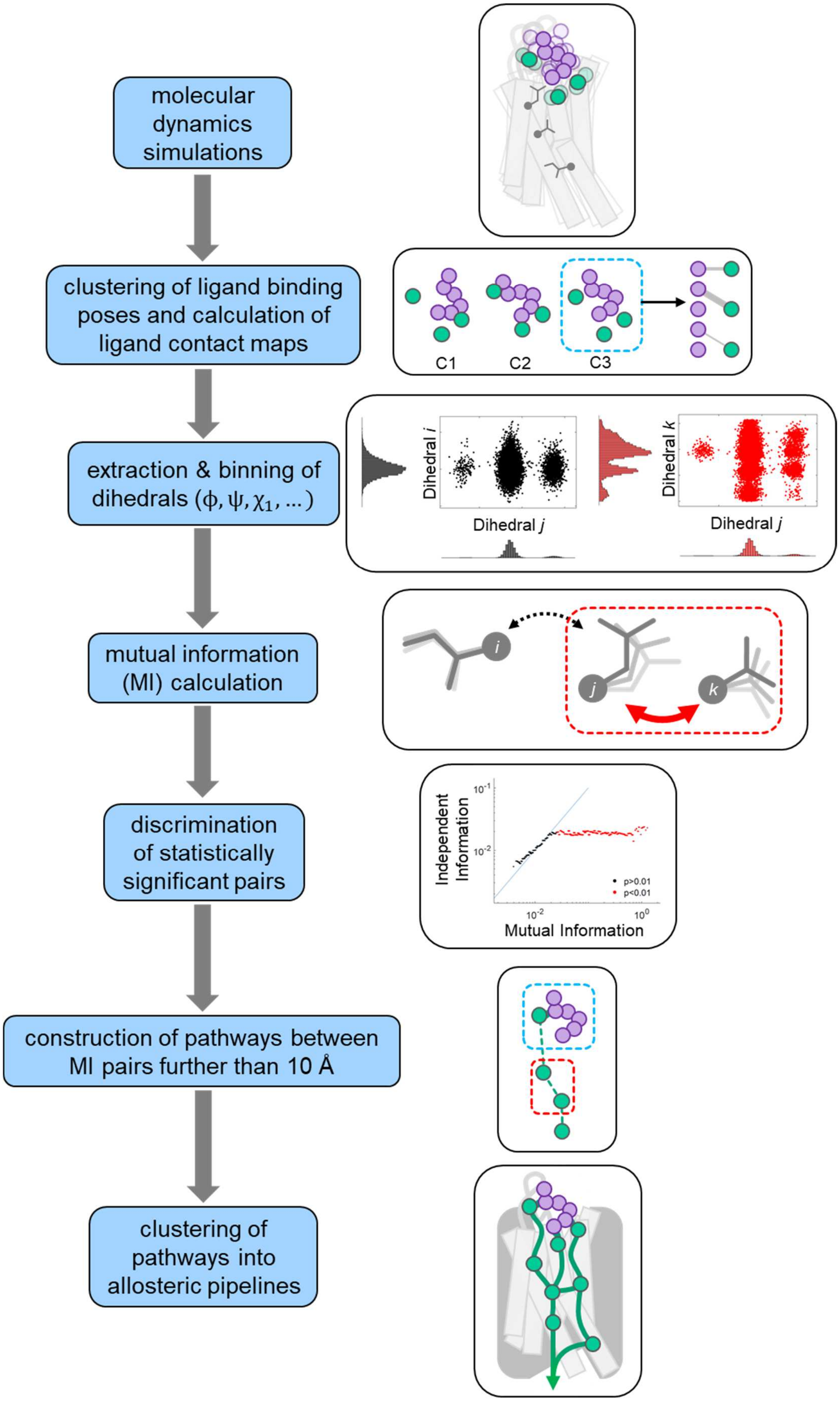
molecular dynamics simulations
clustering of ligand binding poses and calculation of ligand contact maps
C1
C2
C3
extraction & binning of dihedrals ($\phi, \psi, \chi_1$, ...)
Dihedral i
Dihedral j
Dihedral k
Dihedral j
mutual information (MI) calculation
i
j
k
discrimination of statistically significant pairs
Independent Information
Mutual Information
p>0.01
p<0.01
construction of pathways between MI pairs further than 10 Å
clustering of pathways into allosteric pipelines

**Supplementary Table 2. Allosteric strength of transmission hubs across variants.**

Allosteric hubscores calculated from AlloDy for transmission hub residues conserved among WT and designed variants. Hubscores measure the total number of allosteric pathways running through a residue. Since pathways are constructed from pairs of residues exchanging significant amount of mutual information, comparison of hubscores between transmission hubs gives an indication of the relative amount of information passing through these sites (see Methods for detailed calculations). The fractional hubscore among the transmission hubs for each variant is shown in parentheses to enable comparison between variants.

| Hub | F87$^{2.53}$ | L120$^{3.36}$ | H203$^{5.42}$ | W252$^{6.48}$ | N298$^{7.49}$ |
|---|---|---|---|---|---|
| WT | 670 (0.27) | 792 (0.32) | 310 (0.12) | 517 (0.21) | 220 (0.09) |
| Csel2:Y7L | 621 (0.22) | 378 (0.13) | 482 (0.17) | 338 (0.12) | 1017 (0.36) |
| Cdyn:V3Y | 302 (0.14) | 648 (0.29) | 395 (0.18) | 156 (0.07) | 733 (0.33) |
| Csedy:V3Y-Y7L | 881 (0.21) | 1422 (0.34) | 772 (0.18) | 171 (0.04) | 990 (0.23) |

**Supplementary Figure 7. Predicted allosteric couplings in the WT CXCR4:WT CXCL12 complex. (A-C)**. Predicted allosteric pipelines (solid lines) starting from the peptide and running through the receptor towards the intracellular side are calculated for substates C1 (**A**), C2 (**B**), and C3 (**C**) and represented schematically as follows using 3 layers of residues from left to right. Left layer: Peptide residues shown as grey spheres. Middle layer: Receptor residues in the extracellular peptide binding pocket shown as ovals. Those allosterically coupled to peptide residues (connected by a solid line) are defined as allosteric triggers. Right layer: Allosteric transmitter residues coupled to allosteric triggers shown as ovals in the receptor core and located in distinct transmembrane helices (TM 2,3,5,6,7). Receptor residues are colored according to their level of sequence conservation in CXCR4.

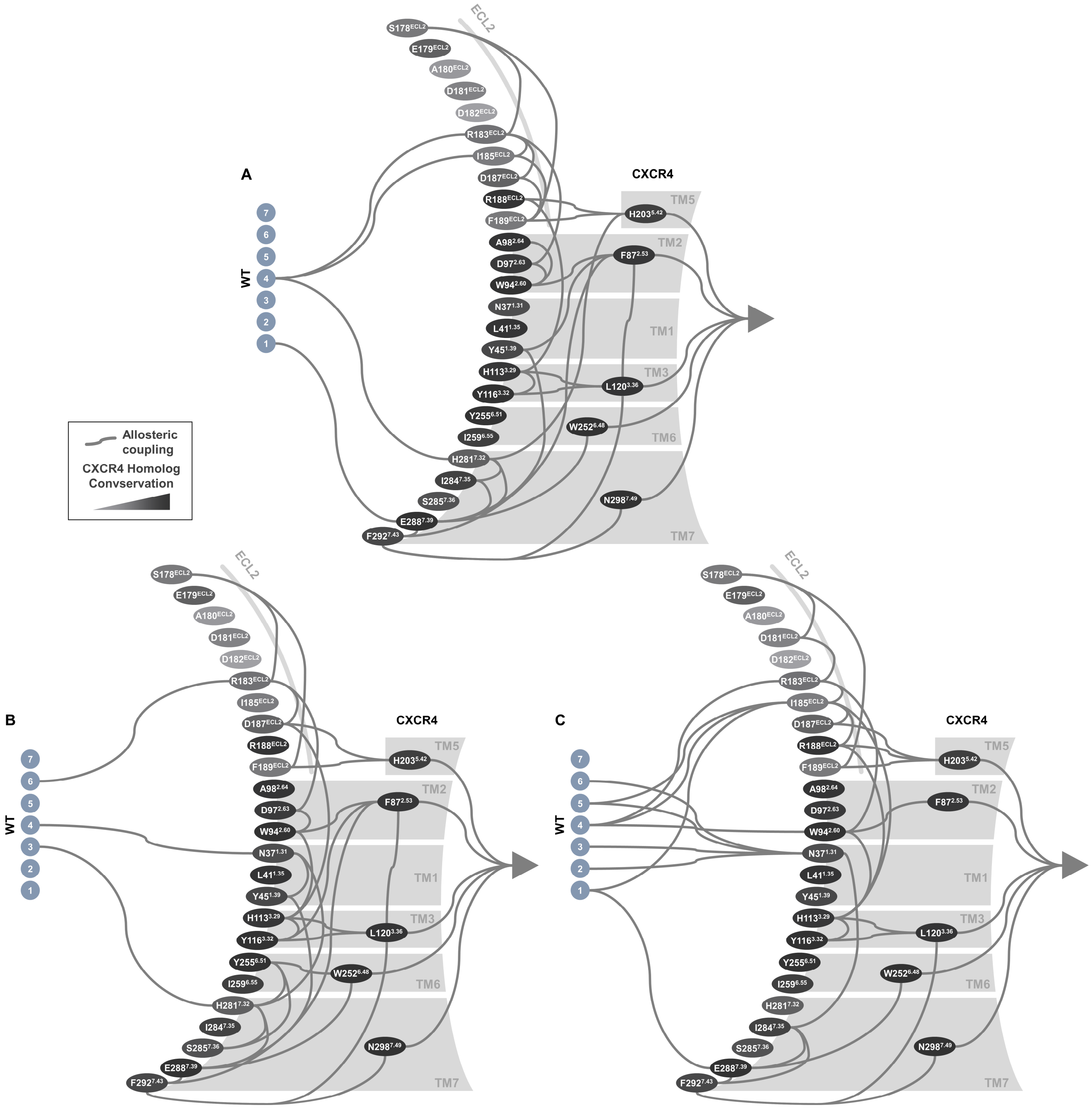
A
B
C
CXCR4
WT
1
2
3
4
5
6
7
ECL2
TM1
TM2
TM3
TM5
TM6
TM7
S178ECL2
E179ECL2
A180ECL2
D181ECL2
D182ECL2
R183ECL2
I185ECL2
D187ECL2
R188ECL2
F189ECL2
A98 2.64
D97 2.63
W94 2.60
N37 1.31
L41 1.35
Y45 1.39
H113 3.29
Y116 3.32
Y255 6.51
I259 6.55
H281 7.32
I284 7.35
S285 7.36
E288 7.39
F292 7.43
H203 5.42
F87 2.53
L120 3.36
W252 6.48
N298 7.49
Allosteric coupling
CXCR4 Homolog Convservation

**Supplementary Figure 8. Predicted allosteric couplings in the CseI2:Y7L CXCL12 complex. (A-C)**. Predicted allosteric pipelines (solid lines) starting from the peptide and running through the receptor towards the intracellular side are calculated for substates C1 (**A**), C2 (**B**), and C3 (**C**) and represented schematically as follows using 3 layers of residues from left to right. Left layer: Peptide residues shown as grey spheres. Middle layer: Receptor residues in the extracellular peptide binding pocket shown as ovals. Those allosterically coupled to peptide residues (connected by a solid line) are defined as allosteric triggers. Right layer: Allosteric transmitter residues coupled to allosteric triggers shown as ovals in the receptor core and located in distinct transmembrane helices (TM 2,3,5,6,7). Receptor residues are colored according to their level of sequence conservation in CXCR4. Mutated peptide and receptor residues are colored in red.

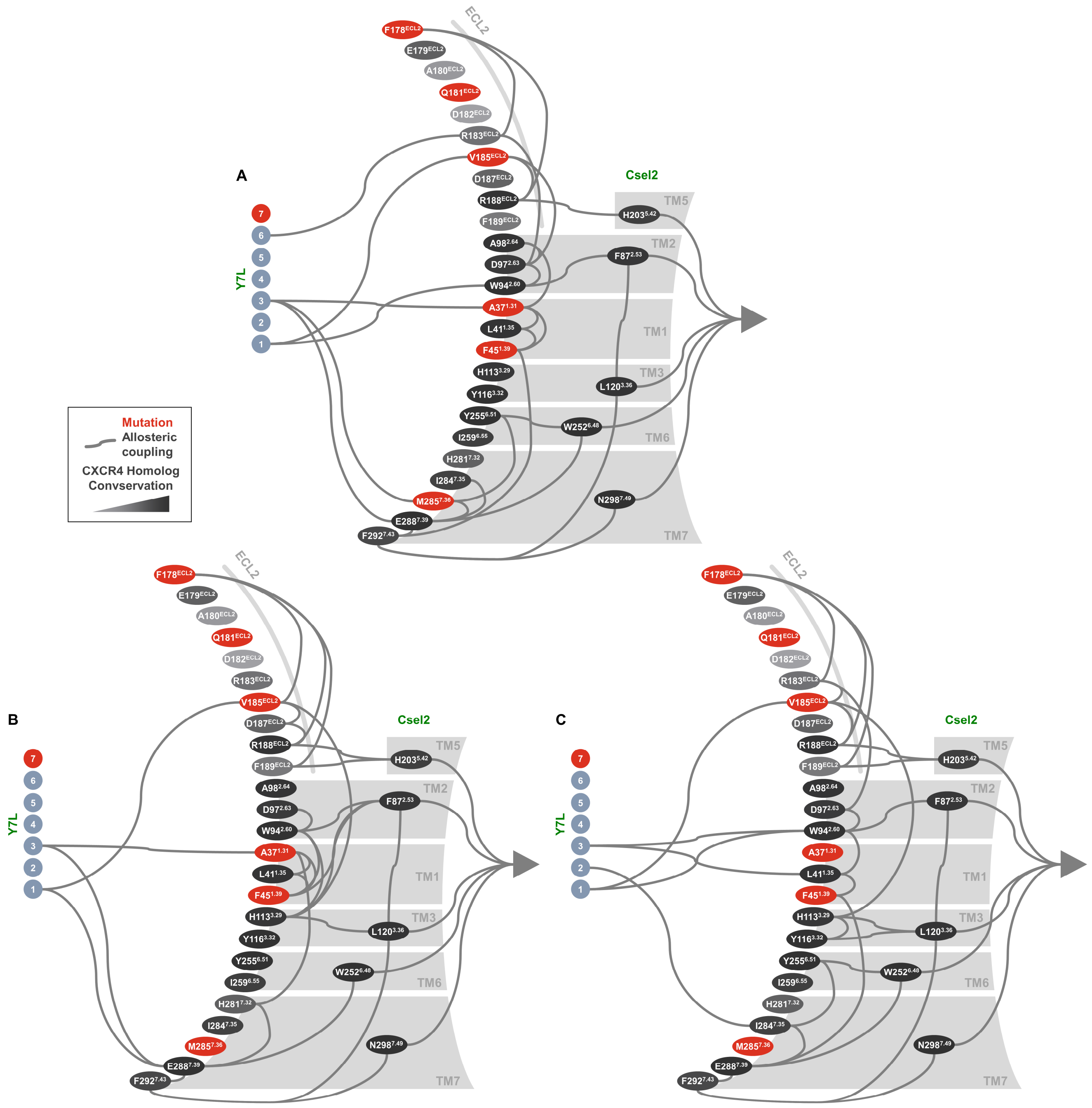
A
B
C
ECL2
Csel2
Y7L
7
6
5
4
3
2
1
F178ECL2
E179ECL2
A180ECL2
Q181ECL2
D182ECL2
R183ECL2
V185ECL2
D187ECL2
R188ECL2
F189ECL2
A98 2.64
D97 2.63
W94 2.60
A37 1.31
L41 1.35
F45 1.39
H113 3.29
Y116 3.32
Y255 6.51
I259 6.55
H281 7.32
I284 7.35
M285 7.36
E288 7.39
F292 7.43
H203 5.42
F87 2.53
L120 3.36
W252 6.48
N298 7.49
TM5
TM2
TM1
TM3
TM6
TM7
Mutation
Allosteric
coupling
CXCR4 Homolog
Convservation

**Supplementary Figure 9. Predicted allosteric couplings in the Cdyn:V3Y CXCL12 complex. (A-C)**. Predicted allosteric pipelines (solid lines) calculated for substates C1 (**A**), C2 (**B**), and C3 (**C**) are represented schematically as follows using 3 layers of residues from left to right. Left layer: Peptide residues shown as grey spheres. Middle layer: Receptor residues in the extracellular peptide binding pocket shown as ovals. Those allosterically coupled to peptide residues (connected by a solid line) are defined as allosteric triggers. Right layer: Allosteric transmitter residues coupled to allosteric triggers shown as ovals in the receptor core and located in distinct transmembrane helices (TM 2,3,5,6,7). Receptor residues are colored according to their level of sequence conservation in CXCR4. Mutated peptide and receptor residues are colored in red.

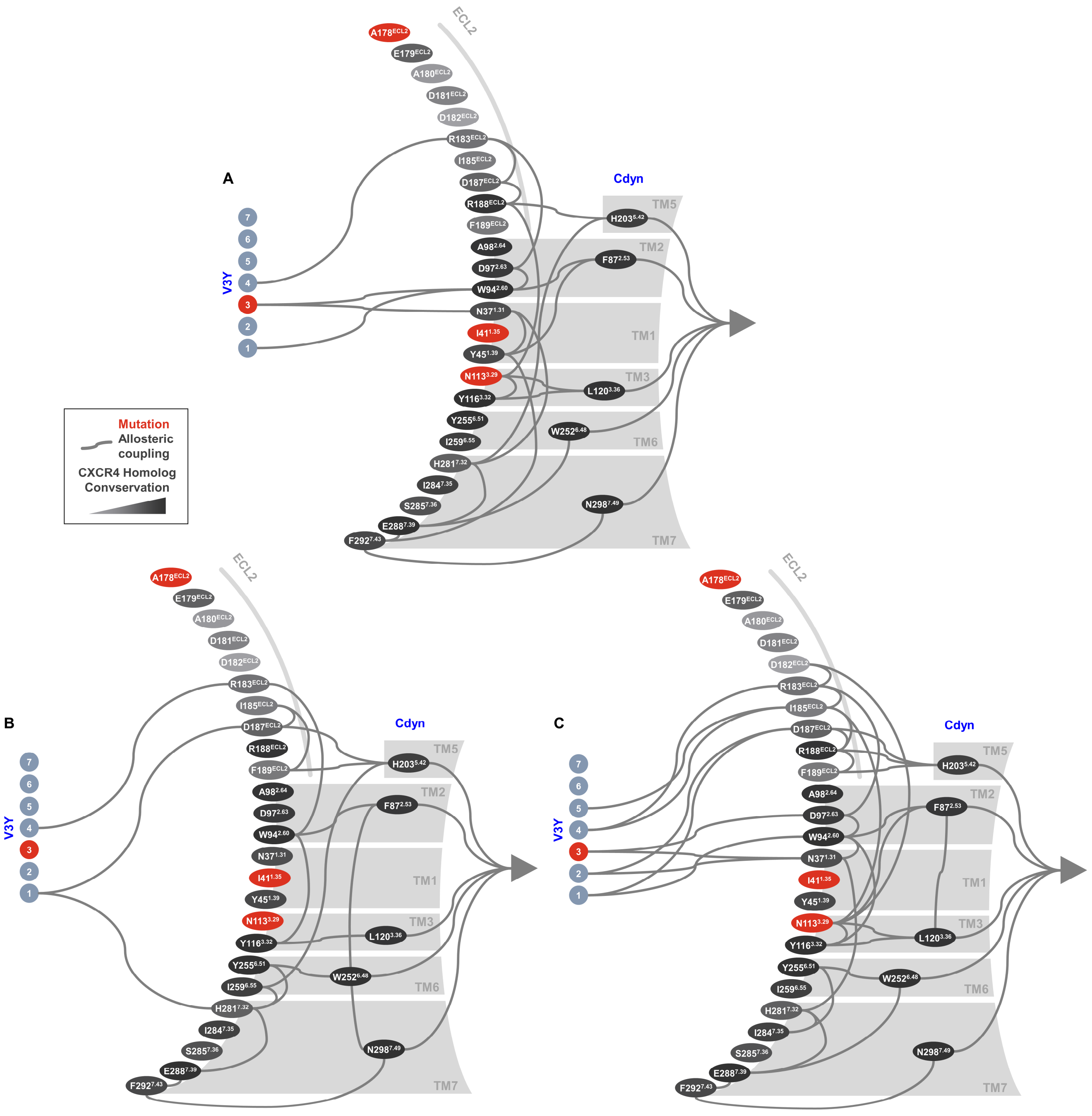
A
B
C
Mutation
Allosteric coupling
CXCR4 Homolog Convservation
V3Y
Cdyn
ECL2
TM1
TM2
TM3
TM5
TM6
TM7
A178ECL2
E179ECL2
A180ECL2
D181ECL2
D182ECL2
R183ECL2
I185ECL2
D187ECL2
R188ECL2
F189ECL2
A98 2.64
D97 2.63
W94 2.60
N37 1.31
I41 1.35
Y45 1.39
N113 3.29
Y116 3.32
Y255 6.51
I259 6.55
H281 7.32
I284 7.35
S285 7.36
E288 7.39
F292 7.43
H203 5.42
F87 2.53
L120 3.36
W252 6.48
N298 7.49

**Supplementary Figure 10. Predicted allosteric couplings in the Csedy:V3Y-Y7L CXCL12 complex. (A-C)**. Predicted allosteric pipelines (solid lines) starting from the peptide and running through the receptor towards the intracellular side are calculated for substates C1 (**A**), C2 (**B**), and C3 (**C**) and represented schematically as follows using 3 layers of residues from left to right. Left layer: Peptide residues shown as grey spheres. Middle layer: Receptor residues in the extracellular peptide binding pocket shown as ovals. Those allosterically coupled to peptide residues (connected by a solid line) are defined as allosteric triggers. Right layer: Allosteric transmitter residues coupled to allosteric triggers shown as ovals in the receptor core and located in distinct transmembrane helices (TM 2,3,5,6,7). Receptor residues are colored according to their level of sequence conservation in CXCR4. Mutated peptide and receptor residues are colored in red.

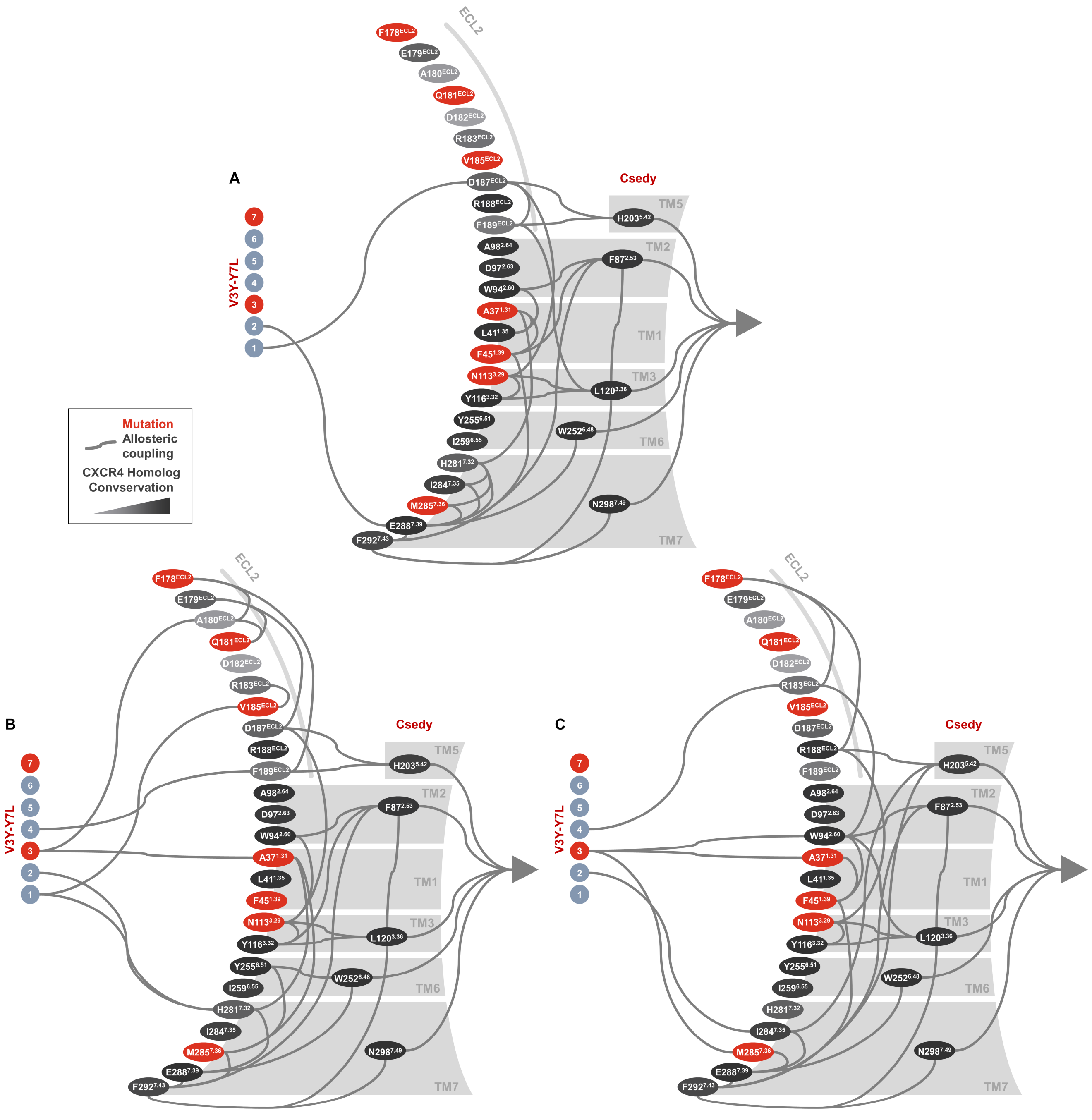
A
B
C
ECL2
Csedy
V3Y-Y7L
7
6
5
4
3
2
1
TM5
TM2
TM1
TM3
TM6
TM7
F178ECL2
E179ECL2
A180ECL2
Q181ECL2
D182ECL2
R183ECL2
V185ECL2
D187ECL2
R188ECL2
F189ECL2
A98 2.64
D97 2.63
W94 2.60
A37 1.31
L41 1.35
F45 1.39
N113 3.29
Y116 3.32
Y255 6.51
I259 6.55
H281 7.32
I284 7.35
M285 7.36
E288 7.39
F292 7.43
H203 5.42
F87 2.53
L120 3.36
W252 6.48
N298 7.49
Mutation
Allosteric coupling
CXCR4 Homolog Convservation

**Supplementary Table 3. Equilibration restraints used for molecular dynamics simulations of receptor—peptide complexes.** Timing and position restraints used for all simulated systems during equilibration in 6 steps.

| Step | 1 | 2 | 3 | 4 | 5 | 6 |
|---|---|---|---|---|---|---|
| Time [ps] | 125 | 125 | 125 | 250 | 250 | 250 |
| Timestep [fs] | 1 | 1 | 1 | 2 | 2 | 2 |
| Ensemble | NVT | NVT | NPT | NPT | NPT | NPT |
| Position restraints [kJ/(mol·nm)] | | | | | | |
| Backbone | 4000 | 2000 | 1000 | 500 | 200 | 50 |
| Side-chain | 2000 | 1000 | 500 | 200 | 50 | 0 |
| Lipid & Ligand | 1000 | 400 | 400 | 200 | 40 | 0 |
| Dihedral | 1000 | 400 | 200 | 200 | 100 | 0 |

**Supplementary Table 4. Simulated time of each system.** Number of runs and simulated time for each run and total simulation time for each receptor—peptide system.

| System (Receptor:Peptide) | Number of Runs | Simulated Time of Every Run | Total Simulated Time |
|---|---|---|---|
| WT:WT | 7 | 3 x 200 ns<br>4 x 300 ns | 1.8 μs |
| Cdyn:V3Y | 7 | 2 x 200 ns<br>5 x 300 ns | 1.9 μs |
| Csedy:V3Y-Y7L | 7 | 3 x 200 ns<br>4 x 300 ns | 1.8 μs |
| Csel2:Y7L | 5 | 5 x 300 ns | 1.5 μs |
| CCR5:RANTES | 5 | 5 x 300 ns | 1.5 μs |

**Supplementary Table 5. Variability of each system explained by principle component analysis.**

Variability explained by the first and second principle components (PC1, PC2) for each receptor—peptide system.

| System (Receptor:Peptide) | Variability explained by PC1 (%) | Variability explained by PC2 (%) |
|---|---|---|
| WT:WT | 59.95 | 16.75 |
| Cdyn:V3Y | 67.47 | 14.72 |
| Csedy:V3Y-Y7L | 52.37 | 21.34 |
| Csel2:Y7L | 28.33 | 20.52 |
| CCR5:RANTES | 53.66 | 19.09 |
